# The tooth on-a-chip: a microphysiologic model system mimicking the pulp-dentin interface and its interaction with biomaterials

**DOI:** 10.1101/748053

**Authors:** Cristiane Miranda França, Anthony Tahayeri, Nara Sousa Rodrigues, Shirin Ferdosian, Regina Puppin-Rontani, Jack L. Ferracane, Luiz E. Bertassoni

**Affiliations:** Department of Restorative Dentistry, School of Dentistry, Oregon Health & Science University, Portland, OR, USA; Post-Graduation Program in Dentistry, Federal University of Ceará, Fortaleza, Ceará, Brazil; School of Dentistry, University of Campinas, Piracicaba, Sao Paulo, Brazil; Center for Regenerative Medicine, School of Medicine, Oregon Health & Science University, Portland, OR, USA; Department of Biomedical Engineering, School of Medicine, Oregon Health & Science University, Portland, OR, USA; Cancer Early Detection Advanced Research Center (CEDAR), Knight Cancer Institute, Portland, OR, USA

**Keywords:** stem cells, bioengineering, MMPs, biocompatibility, restorative materials, odontoblast

## Abstract

The tooth has a unique configuration with respect to biomaterials that are used for its treatment. Cells inside of the dental pulp interface indirectly with biomaterials via a calcified permeable membrane, formed by a dentin barrier which is composed of several thousands of dentinal tubules (~2 µm in diameter) connecting the dental pulp tissue to the outer surface of the tooth. Although the cytotoxic response of the dental pulp to biomaterials has been extensively studied, there is a shortage of in vitro model systems that mimic the dentin-pulp interface, enabling an improved understanding of the morphologic, metabolic and functional influence of biomaterials on live dental pulp cells. To address this shortage, here we developed an organ-on-a-chip model system which integrates cells cultured directly on a dentin wall within a microdevice which replicates some of the architecture and dynamics of the dentin-pulp interface. The tooth-on-a-chip is made out of molded polydimethylsiloxane (PDMS) with a design consisting of two chambers separated by a dentin fragment. To characterize pulp cell responses to dental materials on-chip, stem cell-derived odontoblasts were seeded onto the dentin surface, and observed using live-cell microscopy. Standard dental materials used clinically (2-hydroxyethylmethacrylate - HEMA, Phosphoric Acid - PA, and Adper-Scotchbond - SB) were tested for cytotoxicity, cell morphology and metabolic activity on-chip, and compared against standardized off-chip controls. All dental materials had cytotoxic effects in both on-chip and off-chip systems in the following order: HEMA>SB>PA (p<0.05), and cells presented consistently higher metabolic activity on-chip than off-chip (p<0.05). Furthermore, the tooth-on-a-chip enabled real-time tracking of odontoblast monolayer formation, remodeling, and death in response to biomaterial treatments, and gelatinolytic activity in a model hybrid layer (HL) formed in the microdevice. In conclusion, the tooth-on-a-chip is a novel platform that replicates near-physiologic conditions of the pulp-dentin interface, and enables live-cell imaging to study dental pulp cell response to biomaterials.

## Introduction

Treatment of dental diseases, such as caries or dentin hypersensitivity, require the application of a biomaterial directly onto a cavity formed on the tooth surface. Typically, these involve the attachment of the biomaterial onto dentin – the calcified tissue underlying the outer dental enamel. Therefore, the structural configuration of the tooth results in the formation of an unique interface, where biomaterials contact the dentin matrix, and indirectly allow reaction byproducts and leachates to diffuse into the underlaying dental pulp. Such interactions are enabled by the presence of dentinal tubules (~2 µm in diameter), which are distributed across the dentin matrix, and house odontoblast cell processes that extend a significant length into the dentinal tubules. Therefore, together with the dental pulp, a restored tooth forms an intricate biomaterials/dentin/cell complex (Bertassoni 2017; Bertassoni et al. 2012) that is unique in the body. The biocompatibility and cytotoxicity of dental materials on pulp cells have been extensively studied. Existing model systems include cells cultured on plates (Caldas et al. 2019; Chaves et al. 2012; Schmalz and Galler 2017), larger devices that require specialized equipment, such as the Hume model (Hume 1984), the in-vitro pulp chamber (Hanks et al. 1988), and the dentin barrier test (Schmalz et al. 1999), or ex-vivo models such as the rodent slice culture (Murray et al. 2000), and entire human tooth culture (Camilleri et al. 2014). Despite their extensive usefulness, these models have limited ability for direct observation (i.e. live cell imaging) of the morphologic and metabolic events that occur as cells inside of the tooth become exposed to biomaterials overtime. Arguably, these events hold important information regarding the ability of the tooth to respond to different biomaterials and treatments.

Organs-on-a-chip, integrate microengineered substrates with microfluidics technologies to replicate levels of tissue functionality that are difficult to achieve with conventional 2D or 3D cell culture systems. Similarly, these miniaturized organ systems allow for straightforward experimental control over multifactorial questions that are too difficult to systematically study *in-*vivo (Bhatia and Ingber 2014; Esch et al. 2015). Several organ-on-a-chip models have demonstrated outstanding ability to replicate the complex multicellular architecture, cell-cell/cell-matrix interactions, tissue mechanics, that are naturally present in complex tissues (Huh et al. 2010; Kim et al. 2012; Torisawa et al. 2014). For instance, a recent gut on-chip-model was engineered to replicate the human gut epithelial microvilli, vasculature, microflora, and peristalsis (Kim et al. 2012). Similar methods have also been used to engineer the liver-on-a-chip (Prot et al. 2012), lung-on-a-chip (Huh et al. 2012), bone-marrow-on-a-chip (Torisawa et al. 2014), and several others (Bhatia and Ingber 2014; Esch et al. 2015). Despite these extensive examples, this technology has remained largely underdeveloped in the scope of dental research, and thus the development of an on-chip model of the tooth has remained elusive.

Here we report the development, optimization, and testing of a miniaturized model of the pulp-dentin interface, which we refer to as the ‘tooth-on-a-chip’. We determine the real-time response of pulp-cells to various dental materials such as phosphoric acid, adhesive monomers, and dental adhesives. We then compare the cytotoxicity, metabolic activity, and morphologic changes of an engineered pulp tissue interfaced with the native human dentin as exposed to these materials in comparison to an ISO in-vitro model. We show that the tooth-on-a-chip provides direct visualization of the complexity of the pulp-dentin-biomaterials interface, and enables real-time assessment of the response of pulp cells to dental materials on a level that was was not possible.

## Materials and Methods

### Microdevice fabrication

A master mold was created from a 1 mm-thick sheet of poly-methyl methacrylate (PMMA) (Figure 1A) using a laser cutter. The PMMA molds were used to make an impression on a transparent layer of polydimethylsiloxane (PDMS) (Fig 1B, C, D) which was fabricated with 4 reservoirs for cell medium. Human dentin from third molars extracted for orthodontic reasons according to the institutional ethics committee guidelines was used. Teeth were sectioned into fragments of 500 µm in thickness, inserted into the PDMS device (Figure 1E), and positioned on a plasma-treated coverslip (Figure 1F). The fully assembled microdevice replicates the interface of dentin with the dental pulp on one side and the dental material with dentin on the other, thus forming two accessible chambers representing the “pulp side” and the “cavity side”, respectively (Figure 1G, H). For details, see Supplementary information.

**Figure 1:**
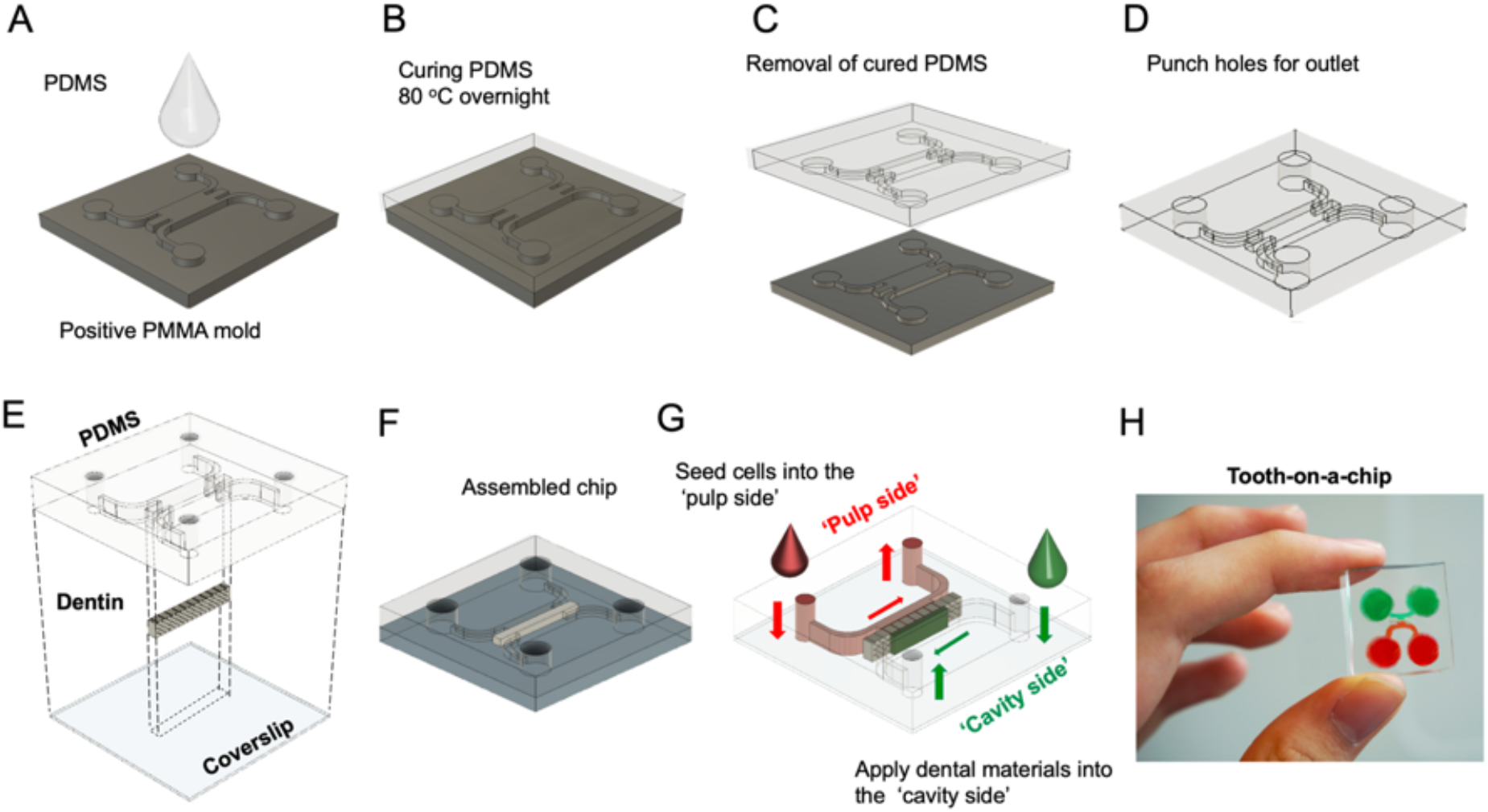
Fabrication of the tooth-on-a-chip. (A) PDMS prepolymer is poured onto a positive PMMA mold, and (B) cured overnight at 80°C. Next, (C) the PDMS is released from the template and (D) 8 mm holes are punched at the end of the channels to form reservoirs. (E) The PDMS and coverslips are plasma-treated, and a dentin fragment is placed in the center, between the two chambers and (F) the system is assembled. Two different chambers representing the ‘pulp side’ and the ‘cavity side’ are formed. (G) The assembled microdevice with dentin as a semipermeable membrane is shown in (H).

### Cell culture

Stem cells from apical papilla (SCAPs) were cultured for 10 days in odontogenic medium, as detailed in the supplementary information. A suspension of 20 µL with 10^5^ SCAP/ml was seeded into the ‘pulp side’ and incubated for 1h to promote cell contact and attachment onto the dentin wall. Next, (Figure 1H) reservoirs were filled with 100 µL of cell medium. Cells were cultured for 7 days with daily cell medium changes.

### Live-cell imaging

To demonstrate that the tooth-on-a-chip enables live-cell imaging of the cells in close contact with the dentin, a monolayer of odontogenically differentiated SCAP cells was imaged overnight, every 30 minutes using a spinning disk confocal microscope (Supplementary information). For the cytotoxicity experiments, on days 1 and 7, chips (n=4) were fixed with 4% paraformaldehyde, stained for actin filaments and nuclei, and imaged using a confocal microscope (LSM 880 Zeiss) (Supplementary information). The whole monolayer in contact with dentin was photographed in 3 consecutive images and analyzed using ImageJ (Fiji, NIH, Maryland, USA).

### Cytotoxicity

After the monolayer formation, three dental materials were tested: (a) HEMA dissolved in cell culture medium at a known cytotoxic concentration of 10 mM, (b) 37% phosphoric acid gel (PA) (Ultradent Products, South Jordan, UT, USA) used to etch the dentin on-chip for 15s, and (c) 35% PA plus Adper Single Bond 2 (SB) (3M/ESPE, St Paul, MN, USA) applied per manufacturer recommendations (Supplementary information). The materials were introduced into the ‘cavity side’ of the dentin, thus forming an interface similar to the dentin-pulp complex in a restored tooth (n=4). Live-cell images of SCAPs assembling as a monolayer on the dentin surface were obtained using a spinning disk field scanning confocal microscope (Yokogawa CSU-X1, Japan). To identify dead cells, SCAPs were incubated with 50 nM of Helix NP NIR (Biolegend, San Diego, CA), a DNA-binding dye, diluted in cell culture medium 10 minutes before imaging. This dye does not require rinsing, so cells continued to be imaged to provide a baseline. After three minutes of imaging, 20 mM of HEMA was added to the opposite side of the dentin and live-cell images were taken every 10 minutes for one hour.

We then compared on-chip experiments against experiments performed using the ISO-10993-1 (ISO 2009), which we refer to as the off-chip group. To that end, 10^4^ SCAPs/well were seeded in 96-well plates and after 24 h, cells were supplemented with the ISO recommended concentrations of HEMA, PA, and extracts of SB obtained by immersing the photopolymerized SB disks in culture media (n=6) (Supplementary information). Next, cells were incubated for 24 h with the conditioned medium of each dental material, and the conditioned medium was replaced with the untreated medium for 7 days. Controls consisted of samples cultured in standard culture medium not exposed to the dental materials. Cell metabolic activity was measured using Alamar Blue (ThermoFisher) on days 0, 1, 3, 5, and 7.

### On-chip zymography of hybrid layer degradation

After determining the cytotoxicity of HEMA, PA, and SB, we evaluated the contribution of MMPs released by dental pulp cells in the degradation of the hybrid layer (HL) formed after SB treatment. We hypothesized that the immediate response of pulp cells to acid attack and monomer exposure might stimulate cells to secrete substantial amounts of proteases, which may stimulate HL degradation in-vivo – a phenomenon that cannot be measured using traditional off-chip zymography methods without cells. To that end, we applied SB as previously described to the ‘cavity side’ of chips fabricated with or without SCAPs (n=4). On-chip zymography of HL degradation was performed as detailed in supplementary information using fluorescein-conjugated gelatin (DQ™ gelatin, EnzChek Gelatinase Assay Kit, ThermoFisher) in the cavity and pulp sides of the device incubated for at least 48 h, after which the proteolytic activity was imaged via confocal microscopy.

### Statistics

Results were analyzed using either a Student’s t-test, one-way ANOVA or two-way ANOVA followed by Tukey post-hoc tests (α = 0.05) on GraphPad Prism 8.

## Results

### Morphogenesis of SCAP on dentin-pulp interface on-a-chip

The tooth-on-a-chip was fabricated with a 500 µm thick dentin barrier and a monolayer of differentiated SCAPs on the opposite site of the dentin wall to emulate a deep cavity.

Figure 2 A-D shows a time lapse of the monolayer assembly on the dentin wall (Figure 2 A) until the complete attachment to the dentin is visible (Figure 2 D). We also tracked the cell death by incubating SCAPs with Helix NP NIR (Biolegend, San Diego, CA), a fluorescent dye that only binds to the DNA of non-viable cells (Figure 2 E-H). Next, we added 20 mM of 2-hydroxyethyl methacrylate (HEMA) (Esstech, PA, USA) to the ‘cavity side’ of the chip and images were taken every 10 minutes. When cells died, the nuclei emitted fluorescence on the 500-530 nm wavelength. Only cells located up to 60 µm away from the dentin were considered as part of the odontoblast-like monolayer.

**Figure 2:**
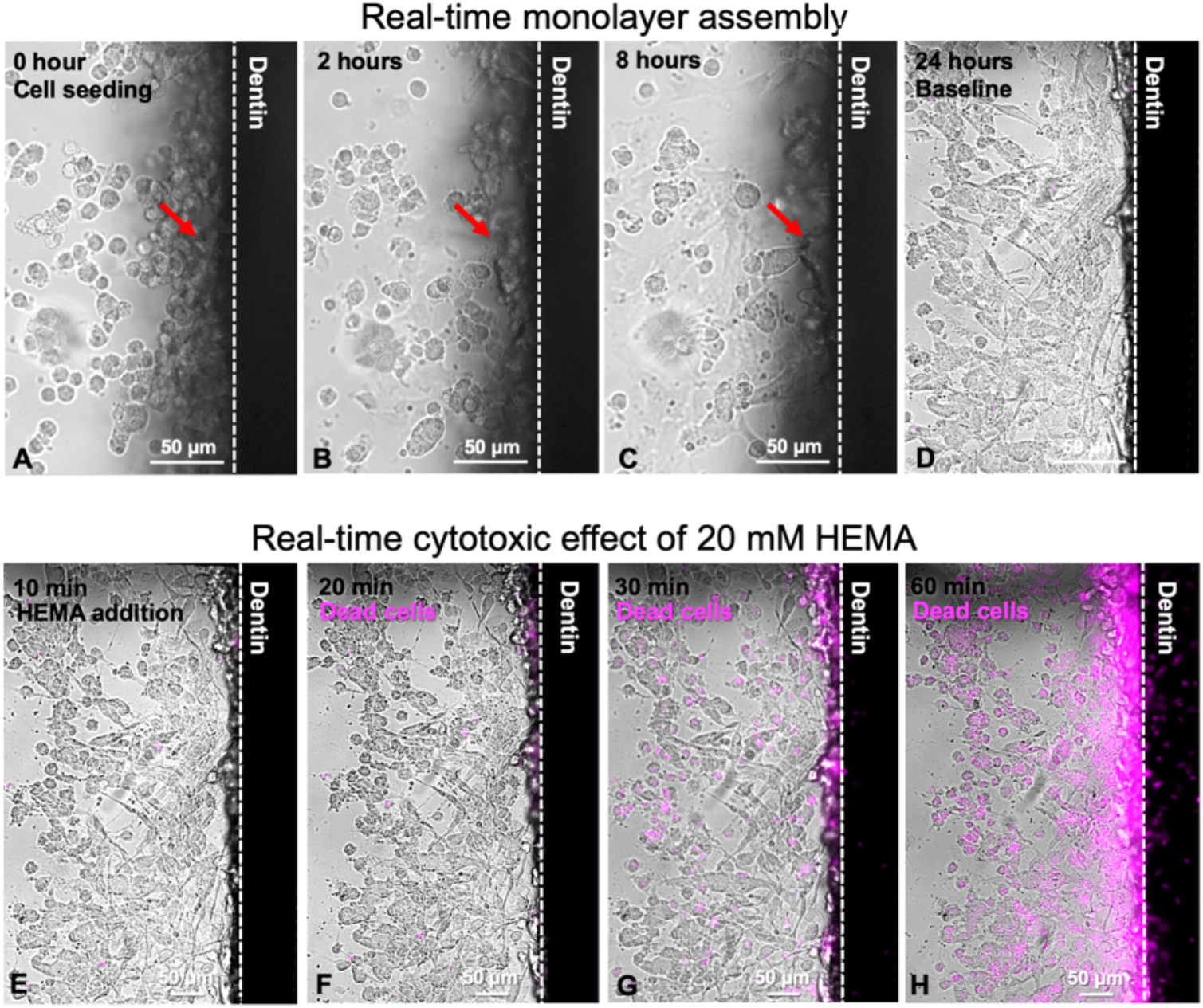
Live-cell imaging on-chip. 10^5^ stem cells from apical papilla were seeded on-chip (A) showing that almost 50% of cells were spread in 4 hours (B) and complete cell spreading was observed in 8 hours (C). After 12 hours the monolayer was completely formed (D). Arrows show the morphological changes of a single cell, which goes from being round and unattached, to a spread morphology as it attaches to the dentin wall after 8 hours (A-D, arrow). Cells cultured on-chip after 24 hours were incubated with a DNA dye (Helix NP NIR) and imaged, demonstrating initial cell viability near 100% (E). Next, 20 mM HEMA was added to the ‘cavity side’ of the chip and after 10 minutes of incubation cells still showed high viability (F). After 30 minutes, almost 50% of cells had their nuclei stained, which is suggestive of high cell death (G). After 60 minutes, nearly all cells were not viable (H). (Confocal microscopy, 647 nm and bright field, dye – Helix NP NIR). Still images extracted from movies 1 and 2 in supplementary material.

While the effects of HEMA were visible on-chip, we also sought to demonstrate the feasibility of real time observation of cell response to biomaterials while the material was being applied. To that end, figure 3 shows a sequence of images of the cell monolayer on-chip as the dentin fragment is exposed to phosphoric acid, following the recommended clinical steps and time of acid application on the tooth. The interaction of the dentin with the acid results in the formation of a number of microbubbles, which appear to stem from the dentin tubules and intertubular dentin matrix (Figure 3 A-D). Interestingly, a discrete contraction of the cell monolayer towards the dentin wall is also seen, as if the cells appeared to contract in response to the acid at the interface with the tooth (Figure 3 E-L).

**Figure 3:**
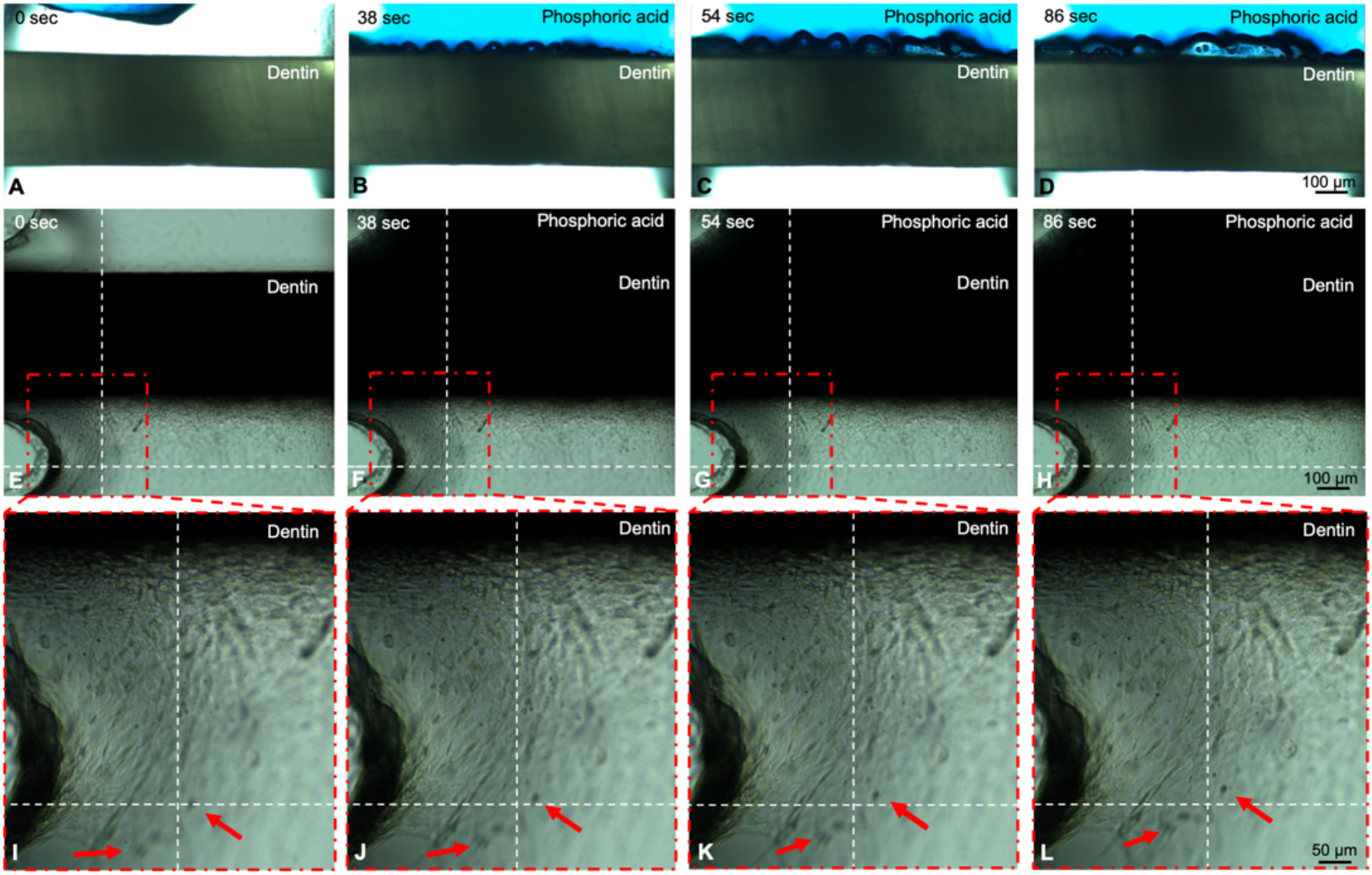
Time lapse showing the process of dentin acid etching with 35% phosphoric acid. Figures A-D show live imaging of the acid reacting with dentin (cells are not seen due to the intensity of light). Figures E-L present the same process with low light intensity to demonstrate the cell monolayer as the dentin is being acid etched. Figures I-L show what appears to the be the contraction of cells in the monolayer as a function of exposure to the phosphoric acid. Red arrows show bundles of cells that appear to move relative to the dotted line, which is provided as reference. Using the scalebar as reference, it is visible that cells appear to move approximately 50 µm over 86 seconds. The supplementary movies (movies 3 and 4) of these events show a clear contraction of the monolayer as a whole.

### Effect of dental materials on SCAP morphology and proliferation

Each tested material elicited apparent cellular injury as early as 24h after treatment for both on-chip (Figure 4 A-D) and off-chip (Figure 4 E-H) samples. The monolayer in the HEMA group consisted of poorly connected round cells with pyknotic nuclei (Figure 4 B) while off-chip HEMA was significantly more cytotoxic, presenting a 10-fold reduction in cell number (p<0.05) relative to the untreated controls (Figure 4 F, I). PA etching of the dentin on-chip caused more discrete monolayer disorganization, cytoplasmic injuries, and separation of cells from the dentin fragment (Figure 4C). Similar cytoplasmic changes were seen in off-chip samples, although overall the effect appeared to be far more significant than off-chip. SB treatment off-chip also caused cytoplasmic changes characterized by a dim actin stain and increased intercellular spaces (Figure 4 D), with a visible decrease in cell number. After 7 days, there were striking differences between on-chip (Figure 4 K-M) and off-chip (Figure 4 O-Q) samples regarding cell number and morphology, except for the untreated control samples (Figure 4 J, N). Quantitatively, cells numbers were significantly higher on-chip compared to off-chip controls after 7 days for all treatments, with both HEMA and SB leading to a significant reduction in cell number relative to untreated controls on day 1 (Figure 4 I-R) (p<0.05).

**Figure 4:**
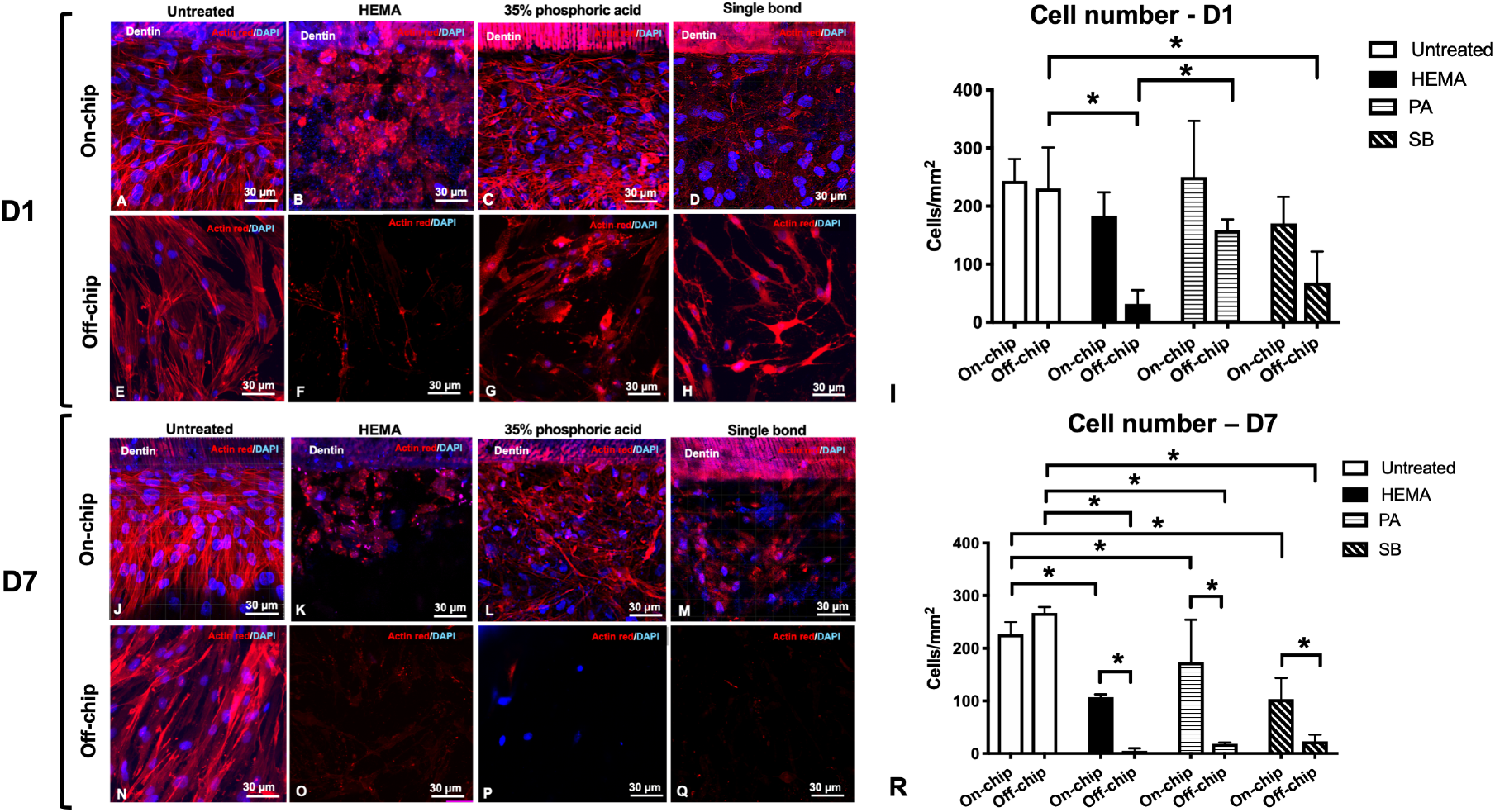
Cytotoxicity of SCAPs on-chip and off-chip. On day 1 untreated samples (A) had SCAP monolayers that were morphologically stable for at least 7 days, while HEMA, PA and SB groups (B, C, and D respectively) showed significant cell morphology changes and decreased cell number on-chip. Cells cultured off-chip on day 1 showed polygonal morphology with oval nuclei in the control group (E), and almost no cells were visible after HEMA treatment. Severe cytoplasmic changes and apparently fragmented nuclei where seen for PA (G) and SB groups (H). Cells on-chip on day 7 showed consistent monolayers on untreated and phosphoric acid samples (J, L) while HEMA and SB groups (K, M) had signs of severe cell degradation. (N) Untreated groups off-chip showed confluent cell monolayers, whereas HEMA, PA, and SB (O, P, Q respectively) displayed very few cells and with faint cytoplasms. Cell count in the monolayer for day 1 and 7 indicated more cells on-chip than off-chip after dental materials application. (Two-way ANOVA, * p < 0.05)

### Metabolic activity

Cells cultured on-chip showed consistently higher metabolic activity than cells cultured off-chip (Figure 5). On-chip, both untreated samples as well as samples treated with PA did not present a significant decrease in metabolic activity over time (Figure 5A, C). Samples treated with HEMA and SB, on the other hand, had a significantly lower metabolic activity only after 7 days (p<0.05). Off-chip, untreated cells presented a spontaneous decrease in cell metabolism as shortly as 24 h (Figure 5A), with HEMA, PA, and SB showing a significant and continuous decrease in metabolic activity from day 1 to day 7 (Figure 5B-D) (p< 0.05).

**Figure 5:**
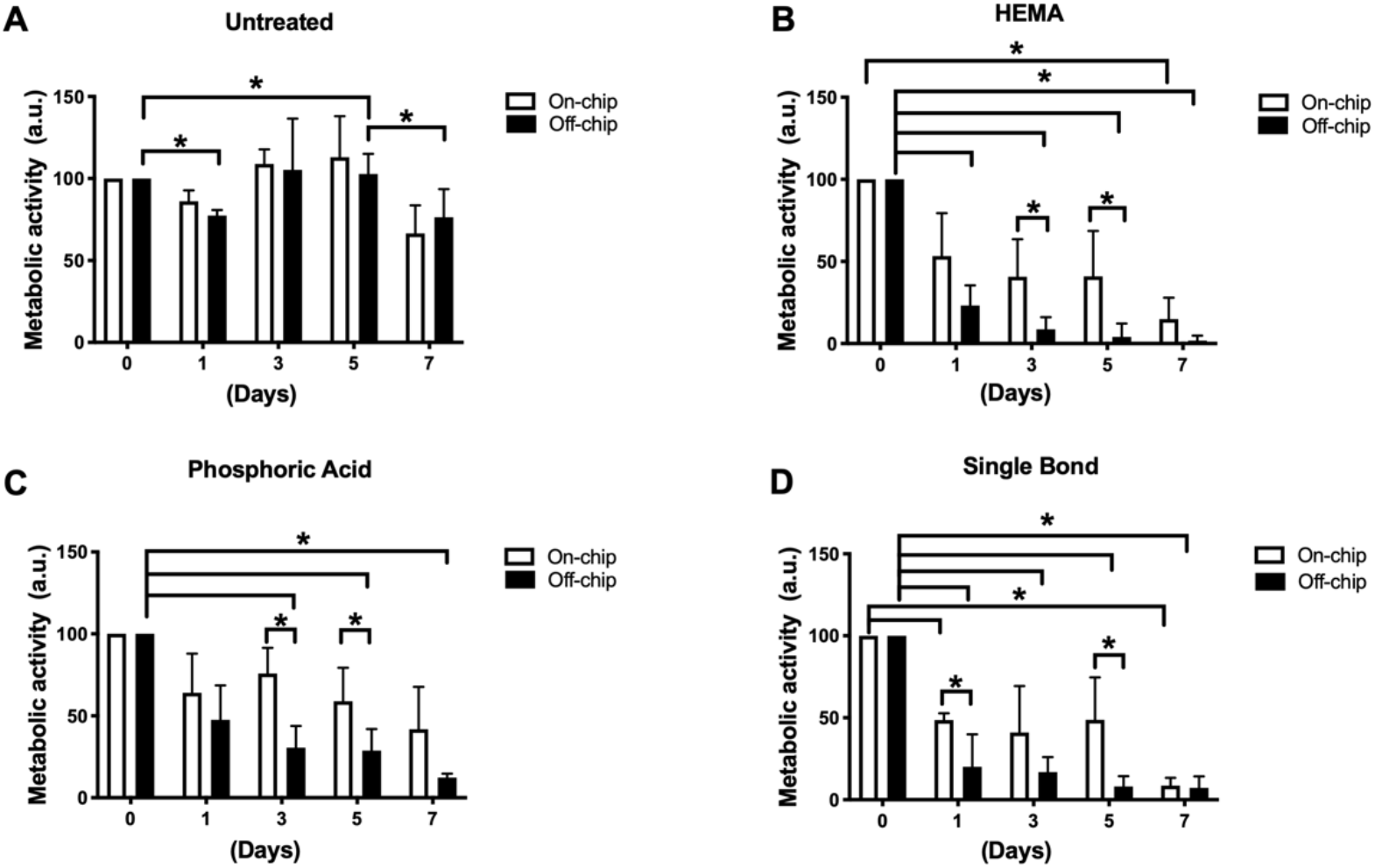
Comparison of metabolic activity between cultures on-chip and off-chip. Cells cultured on-chip had higher metabolic activity than cells cultured off-chip using ISO standards. HEMA (B) was the most cytotoxic material followed by SB (D) and PA (C). (Two-way ANOVA, * p < 0.05)

### In-situ zymography on-chip

To determine the contribution of cellular gelatinases to HL degradation, *in situ* zymography of the HL was performed with fluorescein-conjugated gelatin on chips cultured with or without cells. Green fluorescence, indicative of MMP activity, was visible after 24 h and peaked within 48 h for both groups (Figure 6). Chips seeded with cells had visibly greater HL fluorescence than chips incubated without cells (Figure 6A, F). Interestingly, an intense green fluorescence was detected in colocalization with cell cytoplasm, indicating that the conjugated gelatin was also hydrolyzed at the monolayer (Figure 6H, J). Conversely, chips without cells presented discrete MMP activity at the HL and inside dentin tubules close to the adhesive (Figure 6G).

**Figure 6:**
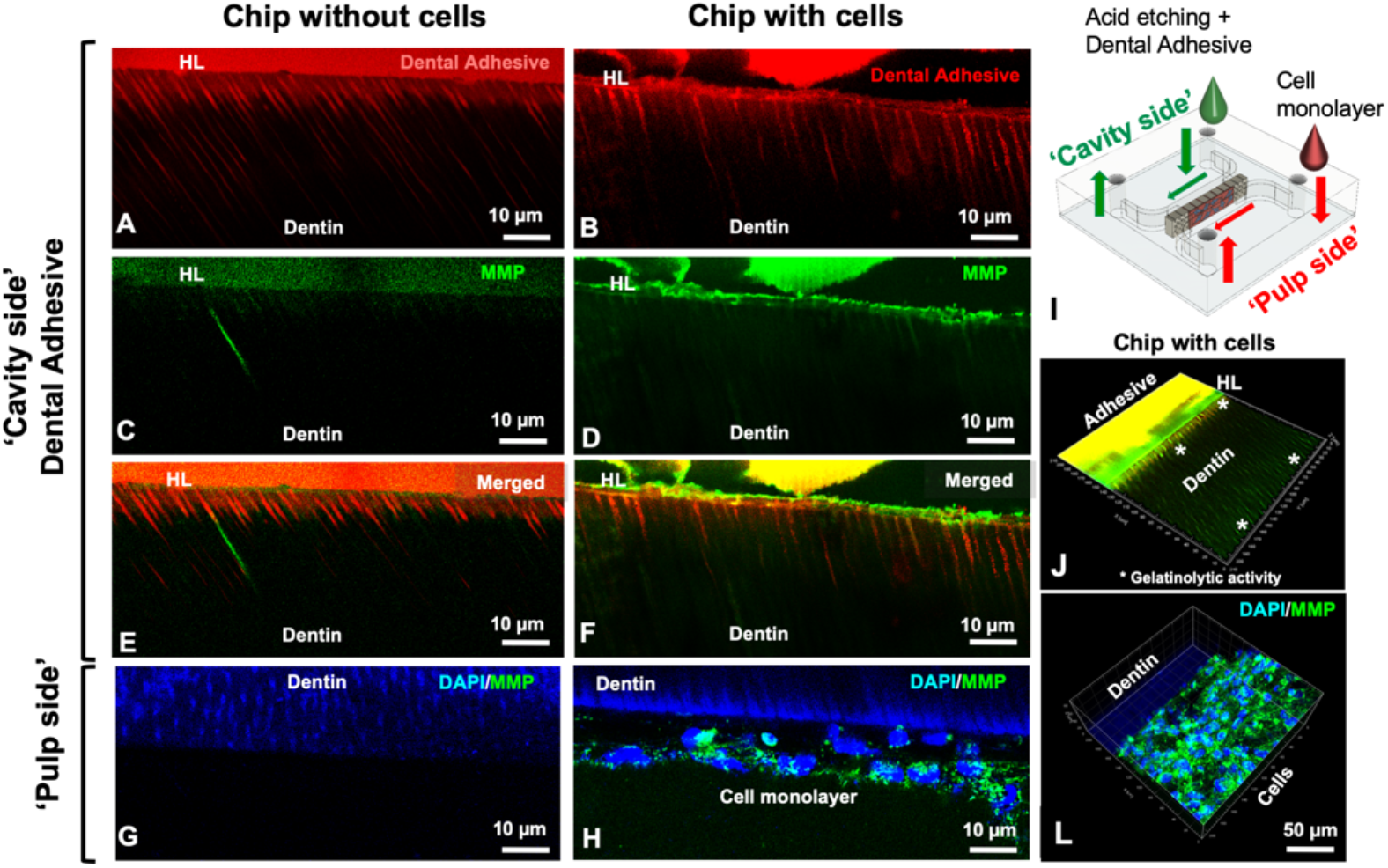
Gelatinolytic activity in the hybrid layer on-chip with and without cells after 48 h. Hybrid layer and tags present in chips without (A) and with cells (B). Fluorescein-conjugated gelatin showing gelatinolytic activity in the HL and dentin tubules (C-F). No evidence of gelatinolytic activity on the ‘pulp side’ for chips without cells (G) while for chips with cells, gelatinolytic activity was co-localized with cell cytoplasm (H). Schematic of the chip (I) and 3D orthogonal view of the adhesive side of a chip with cells (J) showing gelatinolytic activity in the hybrid layer inside dental tubules (*) and on the cell side (L), unquenched gelatin co-localized with cell cytoplasm.

## Discussion

Current understanding of the biological interactions of restorative materials stems from in-vitro experiments using material-based approaches that cannot replicate the complexity of the true biological phenomena occurring at the pulp-dentin interface. Improved model systems that enable direct visualization and quantification of the biological events occurring at the pulp-dentin interface when subjected to different dental materials would address these limitations. To that end, the tooth-on-a-chip provides a highly controllable 3D environment that emulates the pulp-dentin interface in-vivo, allowing for investigation of human pulp cell responses in real-time and in the context in which these interactions occur – that is, at the interface of the dental material with the dentin, forming a permeable barrier to the pulp cells.

One of the key prerequisites for mimicking the dental pulp in-vitro is the establishment of a stable cell-dentin interface comprised of an odontoblast-like cell monolayer that interfaces with the dentin wall and responds to stimuli. First, we tried to encapsulate the cells in a hydrogel to promote a 3D environment on-chip; however, cells did not form monolayers, given that the gels prevent homogenous cell-matrix interactions at the cell-tissue interface. Therefore, we seeded high cell densities on-chip observing complete monolayer formation within 24 hours. Early cell attachment to the dentin was promoted via EDTA pre-treatment, and pre-culturing SCAPs in odontogenic medium for 7-10 days before seeding the cells on-chip. This allowed us to emulate the dentin-pulp interface by fabricating consistent monolayers in a short period of time (Figure 2 A-D), which allowed for easier reproducibility of our experiments and a larger number of replicates. We chose SCAPs because they have a high proliferation rate, potential to differentiate into odontogenic lineage (Miller et al. 2018; Zhang et al. 2015), and to form a typical dentin-pulp like complex in-vivo (Sonoyama et al. 2006). With the tooth-on-a-chip experiments, the combination of odontogenic medium and the dentin matrix provided cells with a microenvironment comprised of biochemical signaling, dentin nanotopography, and stiffness that is favorable to produce monolayers that are stable for several days.

We tested commonly used dental materials on-a-chip and compared their cytotoxic response against well-established ISO controls. Dental materials can interact with the dentin-pulp complex both mechanically, chemically, and indirectly via leachates that travel through the dentin tubules and diffuse through the porosity of the intertubular matrix. Therefore, for comprehensive cytotoxicity screenings, ISO determined various approaches of structuring the material-cell interface, such as direct cell contact, extract test, diffusion tests, dentin barrier and tooth slice models (ISO 2009; ISO/ADA 2008; Schmalz and Galler 2017). A key factor to be considered in the test choice is that dentin has a protective effect on dental pulp cells functioning as a source of growth factors (Ferracane et al. 2013; Salehi et al. 2016) and as a semipermeable barrier, limiting diffusion of leachates to pulp cells (Bertassoni 2017; Bertassoni et al. 2012; Hamid and Hume 1997). Thus, more sophisticated in-vitro models, such as in-vitro pulp chamber, dentin barrier tests, and tooth slices, are preferable (Hanks et al. 1988; ISO/ADA 2008; Schmalz et al. 1999; Schmalz et al. 2001). The tooth-on-a-chip is designed to include a dentin barrier (with a controllable thickness), while being compatible with direct visualization of biological phenomena in real-time, as well as virtually any standard analyses of molecular, metabolic and genetic function, such as PCR, immunostaining and metabolic activity assays like those performed here, as well as others (Bhatia and Ingber 2014). Considerable advantages of this microfluidic devices are the micro-volumes of reagents to maintain the cells in the system - each chip used 100-300 µL of cell culture medium while other systems using dentin disks use more than 1.5 ml per sample (Hanks et al. 1988); the reduced dimensions of the chip, which enable a better use of human tooth since we could get around 10 dentin fragments from each tooth, being able to fabricate 10 chips with only one tooth, and most importantly the ability of real time assessment with extensive experimental control.

On-chip tests with HEMA induced cell death, pyknosis, and cytoplasmic shrinkage within 24 h (Figure 4B, I, R) without completely depleting the pulp-like tissue from viability and response, as it is expected clinically. Cellular responses to HEMA are characterized by an increase in reactive oxygen species, and mitochondrial damage (Bakopoulou et al. 2011; Spagnuolo et al. 2008), which corroborate our morphological findings on-chip. However, off-chip cells were even more sensitive to HEMA, showing a 10-times decrease in cell number and metabolic activity. Potential explanations for this finding, in addition to the lack of dentin barrier function, are that cells cultured off-chip lack the cell-matrix contact with dentin making them more susceptible to external injuries (Schmalz and Galler 2017).

Etching dentin with 35% PA for 15 seconds proved to be harmful to the cell monolayer on-chip, but not enough to fully disrupt it in 24 h. Moreover, after 7 days the monolayer was almost completely reconstituted (Figure 4). Again, this is more consistent with clinical outcomes, where acid-etching elicits the solubilization of dentin matrix components that are important for pulp regeneration (Athirasala et al. 2017; Ferracane et al. 2013; Salehi et al. 2016) instead of causing odontoblast death. Conversely, off-chip experiments showed that the same 15 s of acid application (as recommended by ISO), elicited 90% of cell death after 7 days (Figure 4) and a decrease in metabolic activity (Figure 5), indicating that cells off-chip were far more sensitive to acidic conditions. It seems that results obtained off-chip overestimated PA cytotoxicity.

Additionally, our results suggest that in 24 h the monolayer is partially disrupted, showing visible cytoplasmic changes (Figure 4), while on day 7 the monolayer had disappeared and cell number had decreased to 20-30 cell/mm^2^, indicating the potential toxic effect of leachates over time. Likewise, the on-chip morphological results denote different cellular cytotoxicity mechanisms for HEMA and Single Bond, partially explained by the fact that even though HEMA is a significant component of the Single Bond adhesive composition, HEMA is a freely soluble monomer with a fast action, while Single Bond adhesive is a monomer blend that was photopolymerized, thus releasing potentially toxic leachates much more slowly over time. Cell morphology changes as a consequence of these physical characteristics of the biomaterials, and in the context of the pulp-dentin interface, have remained elusive so far. Additionally, it is possible that other monomers in the composition of Single Bond i.e., UDMA and Bis-GMA may be less soluble than HEMA, in which case they diffuse more slowly (or not at all) through the dentin tubules to affect the cells (Bertassoni 2017; Bertassoni et al. 2012). This same finding could not be reproduced off-chip, and on day 1, Single Bond application proved to be highly cytotoxic to cells, leading to nuclei changes and more than 90% of cell death on day 7.

Lastly, we performed a functional assay with the tooth-on-a-chip, by investigating the role of cellular MMPs in hybrid layer degradation. The release of endogenous dentin MMPs that are bound to collagen fibrils in mineralized dentin and become exposed after acid etching procedures has been implicated in the HL enzymatic degradation, eventually deteriorating the resin-dentin bonding interface (Breschi et al. 2010; Mazzoni et al. 2012; Tjaderhane et al. 2013). Studies conducted with quenched fluorescein-conjugated gelatin in dentin slices or dentin extracts show gelatinolytic activity in the hybrid layer (Breschi et al. 2010; Gu et al. 2018; Mazzoni et al. 2012). However, the cellular role in MMP production has been less explored. In 2009, Lehmann et al. provided strong evidence that self-etching adhesives can stimulate odontoblasts to produce MMP2 (Lehmann et al. 2009). Our system is uniquely capable of culturing cells, and investigating gelatinolytic activity while simultaneously imaging the hybrid layer as formed by adhesive systems. This enables one to assess the collective effect of proteases in HL degradation, including both cell- and matrix-derived proteases, which is more meaningful to the proteolytic activity occurring in-vivo. Here, we fabricated hybrid layers in devices with and without cells, simulating a clinical dental protocol, with all the intermediate steps observed in a clinical setting. Chips with cells presented more gelatinolytic activity in the HL, inside the intratubular dentin, and co-localized with the cell cytoplasm (Figure 6), suggesting that cells actively participate in the degradation of the HL.

Organs-on-a-chip are bioengineering devices that attempt to reconstruct key functions of tissues and organs that cannot be modeled using other existing cell culture systems. The main challenge of an organ-on-chip is to recapitulate in vivo physiology of at least a subset of functions and then to progressively add additional functions over time to speed up the development of new materials, drugs, and disease models (Musah et al. 2017). Similarly to what has been done with other organ-on-a-chip models (Kim et al. 2012; Kim et al. 2016), the tooth on-a-chip opens a wide range of systematic investigations of the dentin-pulp tissue. One of the novelties of the chip is the possibility to treat the dentin using the same protocols used in the dental clinic and to image live-cell morphological changes concomitantly. The time lapses of monolayer formation and disassembling due to HEMA solubilization through dentin tubules, as well as the dentin acid etching and hybrid layer formation, demonstrate the tooth-on-a-chip as a window to track pulp cell cellular and subcellular responses in an environment more consistent with the in-vivo conditions. One limitation of this study, however, is that it is not feasible to cover all aspects of the microfluidic device at once> Future iterations of the device, which are currently being developed, include the addition of controllable flow, built-in biosensors and variable designs for different materials systems. Additionally, key biological functionality will require the addition of functional capillaries, immune cells, and innervation on the ‘pulp side’, and microbiome and salivary flow on the ‘cavity side’. This forms the basis for future studies.

## Author Contribution

CMF and AT contributed equally to this work, carried out the experiments, and analyzed the data. NSR, SF, RPR contributed to the experiments and data analyses. CMF, AT, JLF, and LEB wrote the manuscript. LEB conceived the paper, the chip design, and supervised the project.

## Acknowledgment

We acknowledge Dr. Anibal Diogenes (University of Texas) for the donation of SCAPs. We acknowledge expert technical assistance from Dr. Crystal Chaw in the Advanced Light Microscopy Core at the Jungers Center at Oregon Health & Science University. This project was supported by funding from the National Institute of Dental and Craniofacial Research (R01DE026170 and 3R01DE026170-03S1 to LEB), the Oregon Clinical & Translational Research Institute (OCTRI) - Biomedical Innovation Program (BIP), the Innovation in Oral Care Awards sponsored by GlaxoSmithKline (GSK), International Association for Dental Research (IADR), the Michigan-Pittsburgh-Wyss Resource Center – Regenerative Medicine Resource Center (MPW-RM), the OHSU Fellowship for Diversity and Inclusion in Research (OHSU-OFDIR to CMF).

The authors declare no potential conflicts of interest with respect to the authorship and/or publication of this article.

## Supplementary information

### Fabrication of the microfluidic device

The device design was created with a Computer Aided Design (CAD) software (Autodesk Fusion 360, Autodesk Inc, San Rafael, CA, USA) and positive template was laser cut (Boss LS1416, Boss laser, Sandorf, FL, US) in a polymethylmethacrylate (PMMA) board. Next, the templates were attached to the base of an impression container, molded with PDMS pre-polymer (Figure 1A), an oxygen-permeable, biocompatible polymer, and cured at 80°C overnight (Figure 1B). The set cured-PDMS negative mold was removed from the template (Figure 1C), and had four reservoirs prepared with an 8-mm punch (Figure 1D). The device is comprised of two parallel channels, two perfusable chambers (300 µm W × 1 mm L × 1 mm H) and a central groove that holds a dentin fragment (500 µm W × 1 mm H × 4.5 mm L cut perpendicular to the dentin tubules) (Figure 1 E). PDMS positive mold and coverslip were then plasma cleaned (Plasma Cleaner, PDC-32G, Harrick Plasma, Ithaca, NY, US). PDMS plasma treatment of increases silanol groups (-OH) at the surface of the PDMS so that they form strong covalent bonds (Si– O–Si) when brought together with glass. We did not submit dentin to plasma treatment to prevent any chemical change in its structure. Thus, immediately after plasma treating PDMS mold and coverslips, a dentin fragment was carefully inserted into each PDMS mold using tweezers and the system was assembled onto the glass coverslip using slight pressure, forming a sealed and leak-proof microdevice (Figure 1 F) with two chambers separated by a semi-permeable membrane (dentin) creating distinct microenvironments for each chamber, Figure 1 G,H,I. The dentin fragments had 4.5 mm in length, however the borders were placed within the PDMS holders (Fig 1I), thus the dentin area exposed to dental treatments and cell attachment in the device had about 2-2.5 mm L × 1 mm H and 0,5 mm W.

The microfluidic devices were then sterilized with ultraviolet light (EXFO Acticure 4000, 365 nm, at 8.5 cm distance, light density: 45 mW/cm^2^) for 40 min prior to use. Each tooth on-a-chip was then filled with sterile water until use to prevent dentin dehydration.

### Immunofluorescence

On day 1 and 7, chips from each group (n=4) were rinsed with phosphate-buffered saline (PBS), fixed with 4% paraformaldehyde (v/v) for 1h, rinsed with PBS, permeabilized with 0.1% (w/v) Triton X-100 for 15 min under agitation. Unspecific biding sites were blocked with 1.5 % (w/v) bovine serum albumin (BSA) for 1 h. After washing with PBS, chips were incubated Actin Red 555 (cat. # R37112, Molecular Probes, ThermoFisher) for 1h, rinsed with PBS and incubated with NucBlue (cat. # R37606, Molecular Probes, ThermoFisher) for 30 min at 37 °C. Chips were imaged using a confocal microscope (Zeiss, LSM 880, Germany) with an objective of 20x (Zeiss, Plan-Apochromat 20x/0.8 M-27). The depth of imaging was 100-200 µm, split into at least 20 Z-stacks. Three-dimensional (XYZ) Z-stacks were converted into TIFF files using Zen or Imaris software (v9.1, Bitplane – Oxford Instruments, Zurich, Switzerland).

### Cell culture

Stem cells from apical papilla (SCAPs) (Donation from Anibal Diogenes, University of Texas) were cultured in a Minimal Essential Medium Eagle, alpha modification (⍺ MEM, Gibco, ThermoFisher Scientific, Waltham, USA) supplemented with L-glutamine, 100 U/mL penicillin, 100 µg/mL streptomycin (Sigma-Aldrich), and 10% embryonic stem cell fetal bovine serum (eFBS, ThermoFisher). Culture media was changed every 3 days. Cells were maintained in a humidified incubator (5% CO_2_, 37°C). After reaching 80% of confluency, cells were treated with 0.05% trypsin and passaged to subsequent T75 culture plates. Only cells from passages 3 to 10 were used. Before use in experiments, cells were pre-differentiated for 7-10 days in differentiation medium (10 nM dexamethasone, 10 mM β-glycerophosphate, 50 µg/mL ascorbic acid) (all from Sigma–Aldrich) and 10% eFBS.

Each chip with dentin was treated with 17% ethylenediaminetetraacetic acid (EDTA) for 45 s to promote cell adhesion, and rinsed thoroughly. Next, 20 µL of a 10^5^ SCAP ml^−1^ media suspension was seeded into the pulp chamber and incubated vertically for 1h (37 °C, 100% humidity, 5% CO_2_) for cell attachment onto the dentin walls. Next, the reservoirs were filled with differentiation medium.

### ISO cytotoxicity tests

To validate the chips as a microphysiologic platform to test dental materials, the following samples were used: (a) 2-hydroxyethyl methacrylate (HEMA) (cat. # X9687044, Esstech, PA, USA) dissolved in cell culture medium (10 mM, 0.84% v/v), (b) 37% phosphoric acid gel (PA) (Ultradent Products Inc., South Jordan, UT, USA) dentin etching for 15 s and (c) 35% phosphoric acid dentin etching followed by Adper Single Bond 2 (SB) (cat. #51102, 3M/ESPE, St Paul, MN, USA) application according to the manufacturer recommendation. Briefly, dentin was acid-etched for 15s, rinsed 3 times with distilled water or until complete removal of the acid, dried with absorbent paper cone, then adhesive was applied and light-cured for 20s with a dental light (Valo Ultradent Products Inc, South Jordan, UT, USA). The materials were all introduced to the ‘cavity side’ of the dentin after, thus forming an interface akin to the dentin-pulp interface of a restored tooth (n=4). We then compared on-chip experiments against experiments performed using the International Organization for Standardization (ISO-10993-1) part 5 (ISO 2009). To that end, discs of Adper Single Bond 2 were prepared with 20 µL of the adhesive placed inside cylindrical molds of polydimethylsiloxane (PDMS) (4 mm diameter × 2 mm height) and light-cured for 10 seconds with a Valo Light (Valo Ultradent Products Inc, South Jordan, UT, USA) at a power density of 1650 mW/cm^2^. To assure aseptic conditions, the discs were prepared inside a cell culture hood and measured with a digital caliper immediately after the cure were and then immersed in wells of a 24-well plate filled with 400 µL of SCAPS culture medium for 24 h to keep the same weight/volume proportion of adhesive and liquid as the chip and keeping the ISO recommended range of 0.5-6.0 cm^2^/mL (ISO 2009; ISO/ADA 2008). After the 24 h, the elute was filtered with a 0.22 µm syringe (TPP, Darmstadt, Germany) and stored at 4 °C until use. For HEMA, cells were cultured for 24 h in a 10 mM solution dissolved in cell culture medium. To prepare the phosphoric acid group, 20 µL of the acid gel were dispensed onto a filter paper and immersed into 400 µL of SCAP cell culture medium for 15 s, next the elute was syringe-filtered and stored at 4 °C until use.

SCAPS were seeded in a 96-well plate (10^4^ cells/well) in 200 µL of cell culture medium and after 24h, the culture medium was replaced with the Single Bond, HEMA and phosphoric acid extracts. Untreated cells cultured with SCAP medium served as controls. All groups had n = 6. Cells were incubated for 24h with the extracts and cell culture medium was replaced by regular SCAP medium, and cell cultures were followed for 7 days.

### Inline metabolic activity assay with Alamar Blue

Briefly, 10% (v/v) Alamar Blue reagent (cat. # DAL1025, ThermoFisher) was added to the medium, and chips and 96-well plates were incubated for 18 h to allow viable cells to convert resazurin to resorufin. Subsequently, cell medium with Alamar Blue was collected and read using a microplate reader at 570 nm wavelength absorbance. For the Alamar Blue controls autoclaved Alamar Blue/SCAP medium solution was used as a 100% converted control (control A) and blank Alamar Blue/SCAP medium was used as a non-converted control (control B). To calculate the reduction of viability compared to the negative control, the following equation was used:

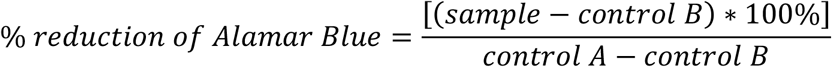

### In-situ zymography of hybrid layer on-chip

To test the hypothesis that SCAPs may contribute to production of proteases and degradation of hybrid layer (HL) in the resin-dentin interface, we prepared the tooth on-a-chip with one group having a monolayer of SCAPs, (as described previously) and the other group with no cells. Dentin was etched for 15s with 35% phosphoric acid gel, rinsed with continuous water irrigation, next, excess water inside the chamber was gently removed with paper cones in order to keep the cell culture medium on the opposite side of the dentin, where cells were seeded. Afterwards 7 µL of Adper Single Bond 2 adhesive labeled with 0.01% rhodamine-isothiocyanate was inserted on the etched dentin, then removed and reinserted in a back and forth movement simulating the two applications recommended in a clinical practice. After 20s the chamber with cells was protected with a photomask and the adhesive was photocured for 20s. DQ™ gelatin conjugated with fluorescein (EnzChek Gelatinase/Collagenase Assay Kit, cat # E-12055, ThermoFisher) was reconstituted in 1 mL of deionized water and the chip reservoirs were filled with 320 µL of cell culture medium and 80 µL of DQ™ gelatin. Chips were incubated at 37 oC, 5% CO2 and 100% humidity. Upon proteolytic digestion, fluorescein-labelled gelatin unquenches yielding highly fluorescent peptides that were live-imaged with a confocal microscope at 48 h.

